# Rat Wetness Response: Sensory Cues, Behavior & Fur-based Drying

**DOI:** 10.1101/2023.09.13.557175

**Authors:** Augustine Triumph Attah, Paola N. Negrón-Moreno, Macarena Amigo-Duran, Linghua Zhang, Max Kenngott, Michael Brecht, Ann M. Clemens

**Affiliations:** Neural Systems & Behavior, Marine Biological Laboratory, 7 MBL Street, Woods Hole, MA 02543 USA; Washington State University, Pullman, USA; Yale University, New Haven, Connecticut, USA; Biomedicine Research Institute of Buenos Aires - CONICET – Partner Institute of the Max Planck Society (IBioBA-MPSP), Argentina; Krieger Mind/Brain Institute, Johns Hopkins University, Baltimore, USA; Brandeis University, Boston, USA; Bernstein Center for Computational Neuroscience, Humboldt University of Berlin, Philippstr. 13 Haus 6, 10115 Berlin, Germany; University of Edinburgh, Simons Initiative for the Developing Brain, 1 George Square, EH8 9JZ, Edinburgh, Scotland, United Kingdom

**Keywords:** Rain, Fur, Drying, Apex, Grooming, Shaking

## Abstract

**It never rains in standard lab-confinements; thus we have limited understanding of animal reactions to water and wetness. To address this issue, we sprayed water on different body parts of rats and measured drying and fur temperature by thermal imaging while manipulating behavior, sensory cues and fur. Spraying water on rats resulted in fur changes (hair clumping, apex formation), grooming, shaking, and scratching. Anesthesia abolished behavioral responses, interfered with fur changes, and slowed drying. Spraying water on different body parts resulted in differential behavioral drying responses. Spraying the head resulted in grooming and shaking responses; water evaporated twice as fast as water sprayed on the animal’s back or belly. We observed no effect of whisker removal on post-water-spraying behavior. In contrast, local anesthesia of dorsal facial skin reduced post-water-spraying behavioral responses. Shaving of head fur drastically enhanced post-water-spraying behaviors, but reduced water loss during drying; indicating that fur promotes evaporation, acting in tandem with behavior to mediate drying. Excised wet fur patches dried and cooled faster than shaved excised wet skin. Water was sucked into distal hair tips, where it evaporated. We propose the wet-fur-heat-pump-hypothesis; fur might extract heat required for drying by cooling ambient air.**

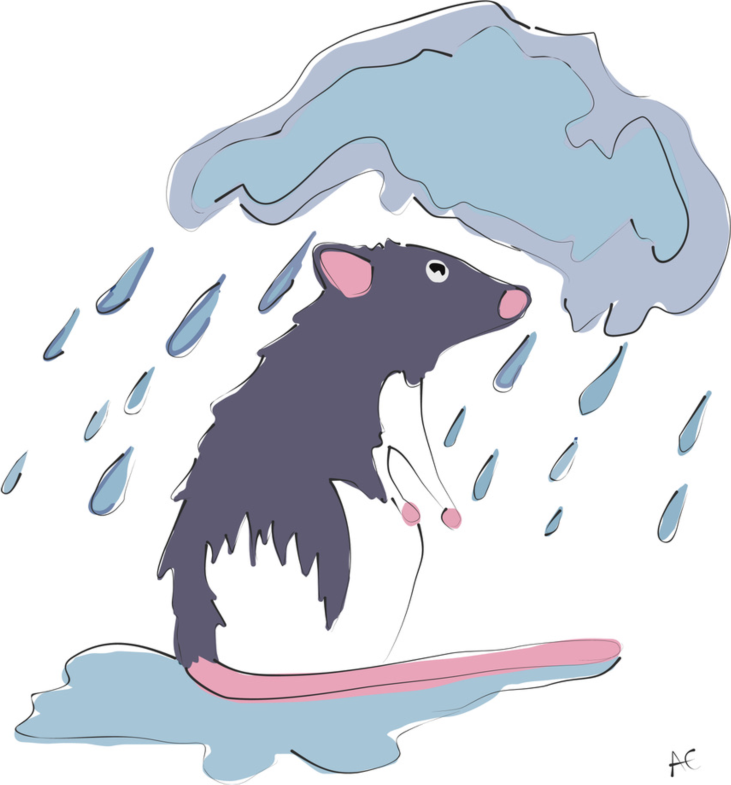

## Introduction

A neuroscientist could easily provide a comprehensive description of what happens in the mammalian brain after a foot-shock or could delineate the cellular and molecular changes that underlie habituation to an air puff to the eye. Fortunately, foot-shocks and air puffs to the eye occur rarely in everyday life. However, neuroscience is often ignorant about the brain’s responses to everyday stimuli. One such everyday-stimulus considered here is rain and the resulting wetness. The relevance of rain and water stimuli is intuitive and important for survival. This process is evidenced by some rodent species adapting to sustain life in wetlands [1], where they frequently need to get wet. Remarkably, water rain stimuli are no less significant in deserts [2]. Yet, we have only limited information about the behavioral and neural reactions to rain and wetness. To address this issue, we sprayed rats with water and characterized behavioral responses, sensory pathways and the drying of the animals.

A particularly striking and well characterized response of land mammals to water is shaking. Such shaking is remarkably efficient in removing water from the wet body. Additionally, fur clumping and fur apexes appear to promote water removal during shaking [3]. Most interestingly, even in plants leaf apex structures appear to contribute to water drainage [4]. Another line of investigation on water stimuli concerns the sensing of wetness. Wetness sensing on the skin is generally considered as a combination of mechanical and thermal responses [5]. In the taste system, water detection is mediated by a group of acid-sensing taste receptor cells [6]. However, in the somatosensory system, sensory pathways dedicated to water sensing *per se* have not been identified.

Thus, we asked the following questions: (1) What behavioral changes result from water spraying in rats? (2) What fur changes result from water spraying in rats? (3) Do behavioral and fur changes depend on the sprayed body part? (4) Which sensory cues contribute to the wetness response? (5) How do behavioral and fur changes impact drying? (6) How does water evaporate from fur? Here we present data suggesting that drying is an active process in rats, with key role of the facial skin in wetness sensing and a major contribution of the apex-forming head fur to drying.

## Results

### Water spraying induces fur changes and stereotyped behaviors in rats

We sprayed rats with water and found that water spraying induced marked changes in behavior and appearance of rats. Specifically, the rat’s fur changed and formed apexes (Figure 1A) as previously described in other mammals [3]. We tested the response to a water-spray challenge in different body parts. After a habituation period to the arena, rats were sprayed either on their head, body, or tail. Post-spray we observed a variety of behavioral changes. Four behaviors stood out, namely whisker grooming (Figure1B), head grooming (Figure1C), body grooming (Figure1D), and shakes (Figure1E). We found that water spraying the head and body significantly increased whisker grooming while spraying the tail had no effect (Figure1F).

**Figure 1.**
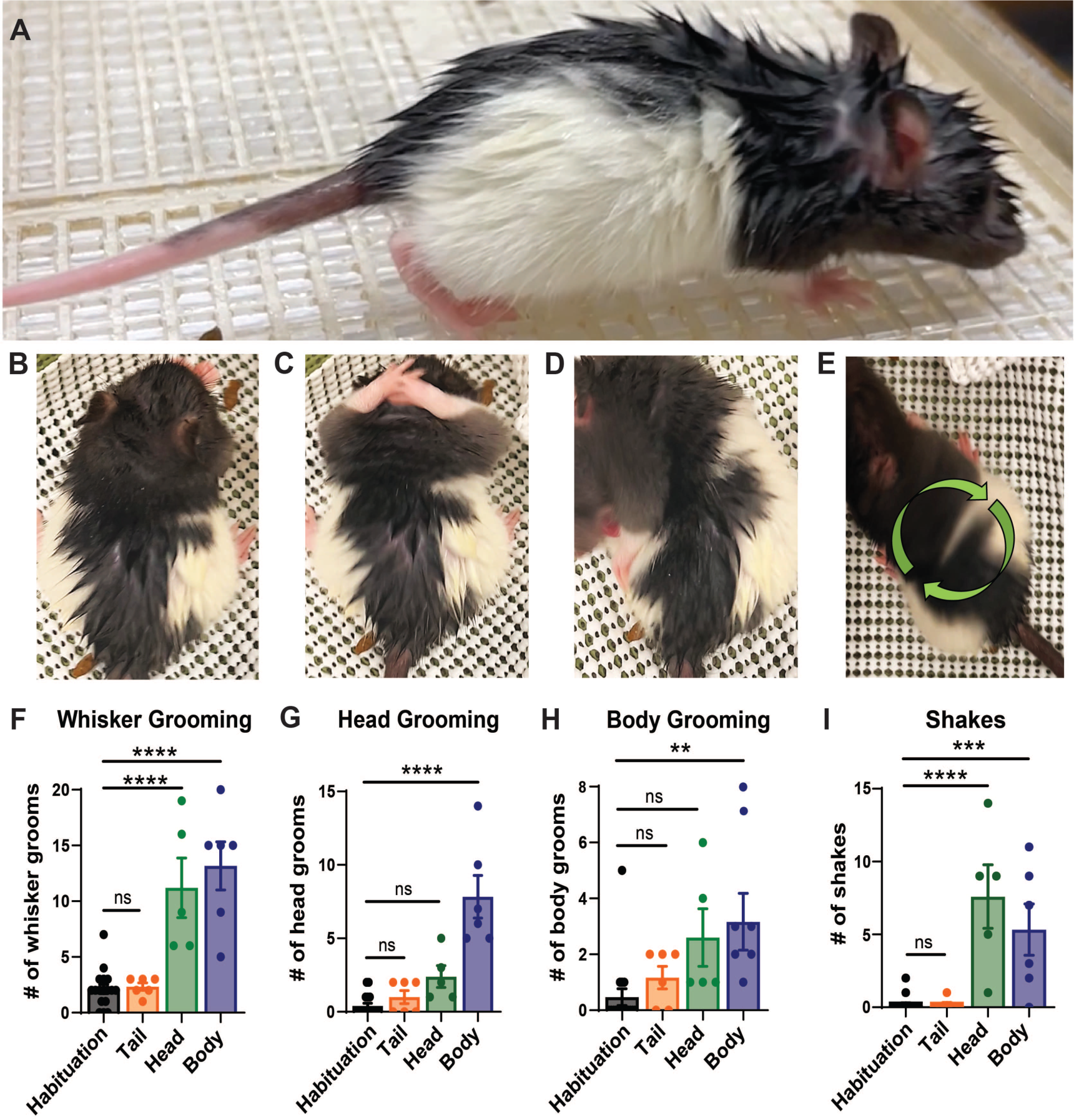
Water spraying induces changes to rat fur and evokes hierarchical and stereotyped behaviors. **(A)** Representative image of a rat after whole-body water spray. **(B-E)** Representative images of stereotyped behaviors induced by wetness (whisker grooming, head grooming, body grooming, and shakes respectively). **(F)** Head-sprayed and body-sprayed rats engage in more whisker grooming initiations than non-sprayed rats. ****p< 0.0001, ****p< 0.0001 respectively. **(G)** Body-sprayed rats engage in more head grooming initiations than non-sprayed rats. ****p<0.0001. Low number of head grooming in head sprayed rats p= 0.1255. **(H)** Body-sprayed rats engage in more body grooming initiations than non-sprayed rats. **p= 0.0086. **(I)** Head-sprayed and body-sprayed rats engage in more shakes than non-sprayed rats. ****p <0.0001, ***p= 0.0010 respectively. Habituation pre-spray: black, n = 17, Tail-sprayed: orange, n=6, Head-sprayed: green, n = 5, Body-sprayed: blue, n = 6. Error bars indicate ± SEM. Data underlying this figure can be found at https://figshare.com/s/8f7b1ef1e553114e20c3

Interestingly, only spraying the body affected the number of head grooming (Figure 1G). One possibility is that the number of head grooming initiations is lower when the head is sprayed (Figure 1G), because the grooming bouts are longer. Further, only spraying the body had a significant effect on body grooming (Figure 1H). Although not significant head spraying showed a tendency toward more body grooming (Figure 1H, p= 0.0729). Shakes were also significantly higher when the head and body received the water spray challenge while spraying the tail had no effect (Figure 1I). These results suggest that different drying strategies are employed depending on where rats are sprayed. However, regardless of spraying location head and body grooming are prioritized. Additionally, these behaviors are stereotyped and are engaged in sequence consistently, albeit to different degrees, when the animal receives a water spray challenge. Our results are consistent with previous studies that have reported shaking [3]. Our observations suggest that water spraying induces behavioral and fur changes, which are stereotyped and partially independent of which body part is wet.

### Anesthesia disrupts fur changes, behavioral wetness responses and drying

The variety and abundance of behaviors led us to ask what the significance of these wetness responses is. To address this question, we sprayed water onto awake and anesthetized animals and measured weight loss as an indicator of drying by placing the animals in a beaker on a scale (Figure 2A).

**Figure 2:**
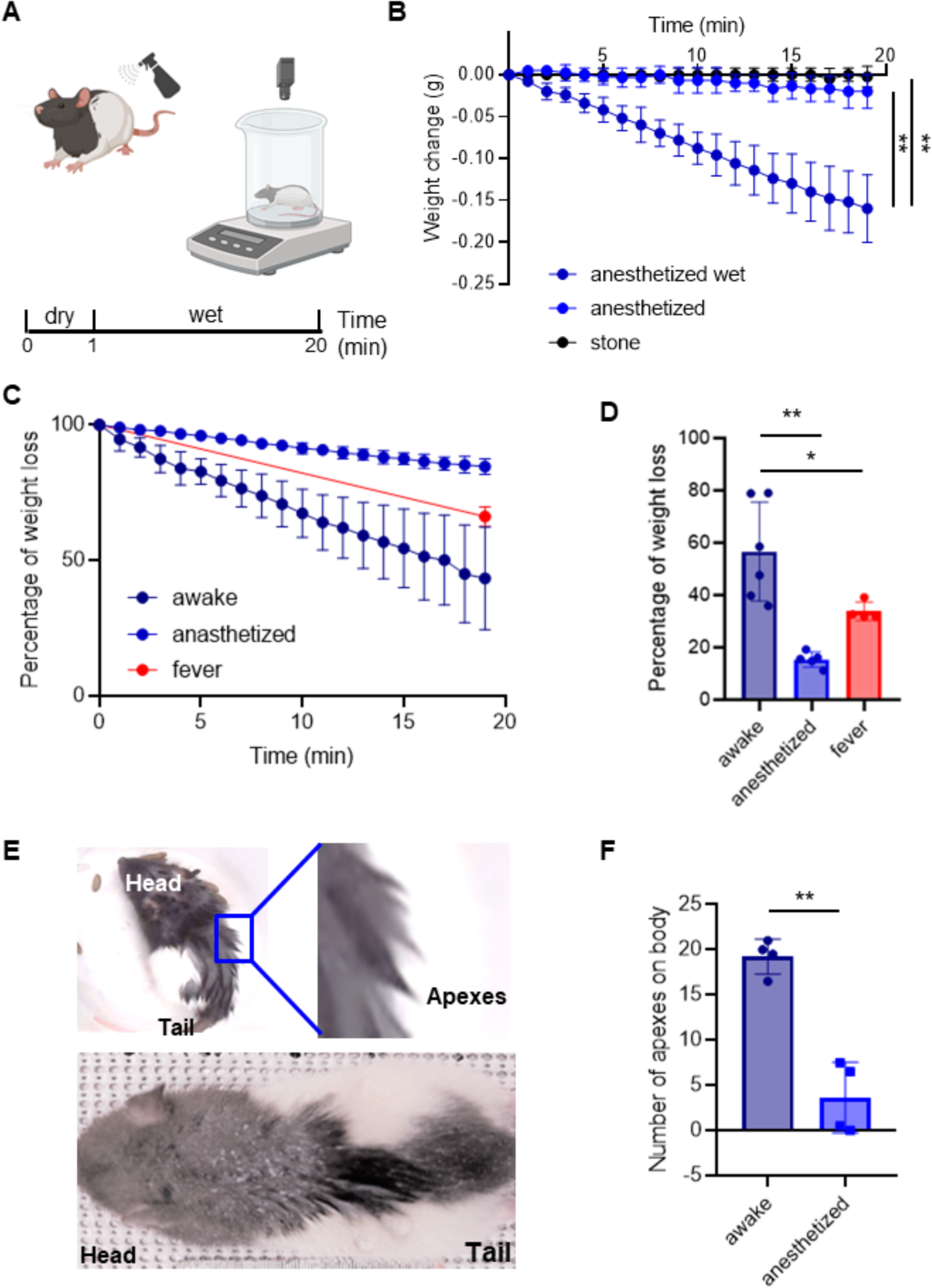
Awake animals dry faster than anesthetized animals. **(A)** Schematic of the weight measurement protocol. Awake and anesthetized animals were weighted, spray five times in the back with DI water and compute the weight loss for five minutes. **(B)** Weight change measurements in stones (n = 4), dry anesthetized (n = 5) and wet anesthetized (n = 6) animals for twenty minutes. At the end of the session, wet animals lose significantly more weight than anesthetized animals and stones (**p=0.001, anesthetized vs anesthetized dry, minute 20; **p=0.001, anesthetized vs stone, minute 20). **(C)** Percentage of water loss in awake (n = 6), anesthetized (n = 5) and anesthetized with artificially increased (by ∼1.5°) body temperature (n = 2, fever) animals for 20 minutes. The total amount of water sprayed on was the weight difference before and after spraying the animal, the weight difference to the weight gain after spraying was measured every minute and plotted in % of the total weight gain by spraying. **(D)** Percentage of water loss in the last minute in awake, anesthetized and fever animals. The median values are significative different (****p<0.0001). Awake animals lose more water than anesthetized (**p=0.0043) and more than fever animals (*p=0.0159). Conventions as in **C**. **(E)** Animals with apex formation after water spray, where the upper one is an awake animal with a zoom showing strong apexes and the bottom one is an anesthetized animal. **(F)** Number of apexes on the fur of the back; values represent average scores of two observers scoring photographs of awake and anesthetized animals five minutes after spraying (*p=0.028). See Supplementary Figure 1 for an alternative method of scoring apexes. Error bars indicate ± SEM. Data underlying this figure can be found at https://figshare.com/s/109204cbfff25915ce87

We first tested the accuracy and sensitivity of our setup to measure drying in a reproducible manner. To this end we weighted stones (n =4) to assess the drift of our measurements over extended time periods and found weight measurements to be stable (Figure 2B). Next, we measured the weight change in anesthetized animals, with and without water being sprayed on (Figure 2B). We found that 20 minutes after spraying wet anesthetized animals changed their weight more than dry anesthetized animals, and the stones, respectively. These data suggest our measurement approach can quantify, to some extent, the evaporation of the water from the bodies of water sprayed animals (Figure 2B). Next, we measured the weight added by the water spraying and compared the percentage of weight that is lost in 20 minutes in awake and anesthetized animals. We found that the awake animals lost more water over 20 minutes than the anesthetized animals (Figure 2C, D). Since anesthesia interferes with homeostatic function it might also have effects on lowering body temperature. We therefore also measured weight loss in anesthetized animals, in which we induced a ∼1.5° higher body temperature (via a heating blanket). We observed that increased body temperature induced higher weight loss in awake animals than in the anesthetized animals (Figure 2D). Additionally, we identified hair clumping and apex formation after water spraying consistent with our previous findings (Figure 1A). To quantify apex formation, two observers independently scored the number of big apexes formed. These scores were collapsed and averaged to compared apex counts between awake and anesthetized animals (Figure 2E). We found that awake animals had more apexes compared to anesthetized animals (Figure 2F).

In addition to subjective scoring of apex numbers, we also developed other means for quantifying apexes (Supplementary Figure S1). To this end, we filtered images of water sprayed animals with an edge detection algorithm. Apexes were apparent as white lines and could then simply quantify apex formation as a total line length measurement over the fur regions. We found that the total length of apex, or fur clumps was significantly decreased at the head region 20 min after spray as a result of drying (n = 3; Figure S1). The disadvantage of our edge detection approach was that the feasibility depended on fur color. Our observations confirm that water spraying alters the fur and suggest that apex formation is amplified during wakefulness, potentially by using behavioral drying strategies. These findings suggests that awake animals dry faster through the incorporation active behavioral strategies and/or fur changes that promote drying.

### Differential drying responses of different body parts after water spraying

We were interested in identifying body parts that were most sensitive to water spraying and how these body parts contributed to drying after exposure to wetness. Since our previous findings suggested that spraying the head and back added the most weight, we decided to focus on characterizing potential differences in the drying process between the head, back and belly of awake rats. To this end, we water sprayed different groups of awake rats on the head, back, or belly (Figure 3A). We immediately placed these rats on a weighing scale and quantified weight loss over five minutes. Awake rats sprayed with water on the head lost weight significantly faster than animals sprayed on the back or belly (Figure 3B, C). We took photographs of the head, back, and belly of awake rats five minutes after exposure to spraying. Rats sprayed with water either on their head, back, or belly displayed robust formation of apexes on their head, back, and belly respectively. However, apexes appeared to be particularly prominent on the head (Figure 3D, E, F). We conclude that the rat’s head dries faster than the back or belly following water exposure. Furthermore, the prominent apexes of the fast-drying head fur point to a role of fur apexes in drying in rats.

**Figure 3.**
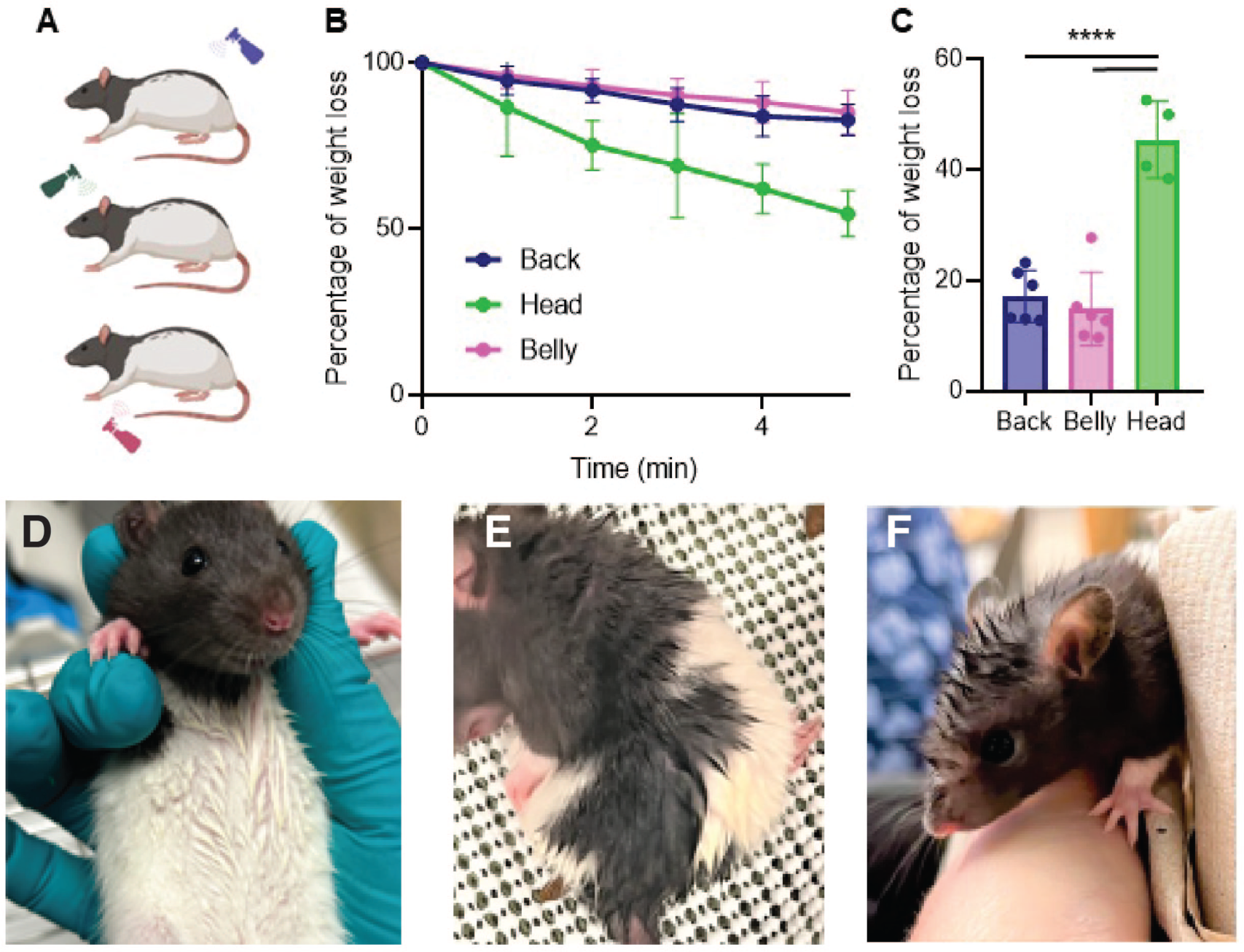
Differential drying responses of different body parts after water spraying. **(A)** Water spraying on the rat’s back (blue bottle, upper), head (green bottle, middle) and belly (pink bottle, lower). **(B)** Drying of different body parts. Percentage of weight loss against time in awake rats sprayed on the back, on the head and on the belly. The total amount of water sprayed on was the weight difference before and after spraying the animal, the weight difference to the weight gain after spraying was measured every minute and plotted in % of the total weight gain. There is a significant difference in weight loss between rats that are sprayed on different parts of the body. **(C)** Percentage of water loss in the last minute in animals sprayed on the back, head and belly. Animals that are sprayed on the head lost more water after 5 minutes than animals that are sprayed on the back (****p<0.0001) and on the belly(****p<0.0001). There is no difference in water loss between animals sprayed on the back and belly (p=0.7945). **(D)** Belly of an awake rat with apexes after water spraying. **(E)** Back of an awake rat with apexes after water spraying. **(F)** Head of an awake rat with apexes after water spraying. Note that head fur apexes are very prominent. Back sprayed: blue, n = 6, head sprayed: green, n = 4, belly sprayed: pink, n = 6. Error bars indicate ± SEM. Data underlying this figure can be found at https://figshare.com/s/109204cbfff25915ce87

### The dorsal facial skin rather than whiskers provides wetness cues

Our initial water spraying experiments suggested that head spraying drives wetness-related behavioral responses. Therefore, the head is likely to be a relevant body part in the drying response. However, how do rats perceive wetness on their head? To determine the role of the whiskers in sensing wetness after trimming (Figure 4A), sprayed these animals, and quantified their behavioral responses post-spray. We found that the sham-trimmed and whisker-trimmed rats displayed the same behavioral responses (Figure 4B). This finding came as a surprise, because whiskers are of great sensory significance to rats and because water spraying induces intense whisker grooming (Figure 1F). This finding suggests that the whiskers do not mediate the wetness response. To establish the sensory pathways activating the wetness response, we than injected one group of rats with 0.5% lidocaine (a powerful local anesthetic) and a second group of rats with saline under the dorsal facial skin of the head (Figure 4C). Consistent with our previous results, behavioral responses are increased after water spraying, however there is a reduced response in animals that received a lidocaine injection in comparison with those that received a saline injection (Figure 4D). Our findings pinpoint the dorsal facial skin as a potential sensor for wetness.

**Figure 4.**
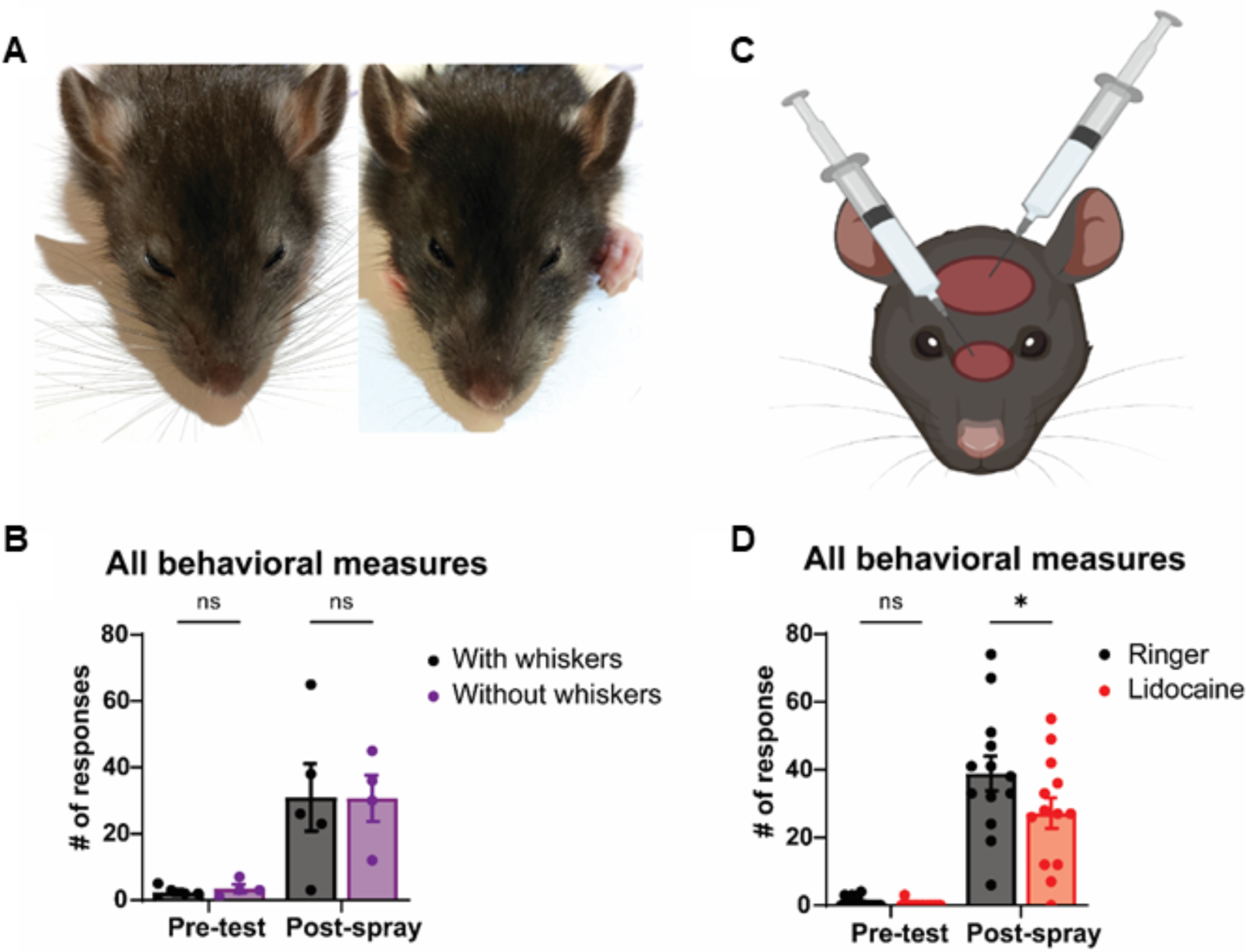
The dorsal facial skin does and whiskers do not drive behavioral responses to head water spraying. **(A)** Representative images of a rat with intact whiskers (left) and with trimmed whiskers (right). **(B)** Whisker trimming does not affect behavioral responses after head spray (p=0.9996). **(C)** Schematic of lidocaine injection sites under the dorsal facial skin. **(D)** Local anesthesia of dorsal facial skin with lidocaine reduces behavioral responses after head spray (*p=0.0384). Ringer: black, n = 13, Lidocaine: purple, n = 13. Intact whiskers: black, n = 5, Trimmed whisker: purple, n = 4. Error bars indicate ± SEM. Data underlying this figure can be found at https://figshare.com/s/2dc01700cbb9c341bfd3

### Shaving the head fur increases behavioral wetness responses but slows drying

Following our finding that whisker trimming does not affect behavioral drying strategies, we tested the role of the head fur in the rat’s wetness response. To this end we shaved the animal’s head (Figure 5A) and compared the drying trajectory and post-spray behavior with that of pseudo-shaved animals. Head-shaved rats performed significantly more drying behaviors than pseudo-shaved rats after head spraying (Figure 5B). Specifically, there is a significant effect on head grooming (Figure 5C) and shakes (Figure 5D). However, there is no effect on whisker grooming (Figure 5E). These findings support our previous observation of fewer behavioral responses after lidocaine injections on the dorsal facial skin. We decided to assess if the increased behavioral response resulted from the increased exposure of the facial skin to water. Interestingly, the most remarkable observation from head-shaved animals concerns the drying of the animals. Paradoxically, considering the increased behavioral responses, we found that shaving the head significantly reduces the rate of drying, evidenced by a reduced rate (percentage) of weight loss across time (Figure 5F). Taken together, our results suggest that head fur plays a critical role in drying, which might be aided by the strong apex formation of head fur. Therefore, the data suggest to us that the head fur might act as water removal cap.

**Figure 5.**
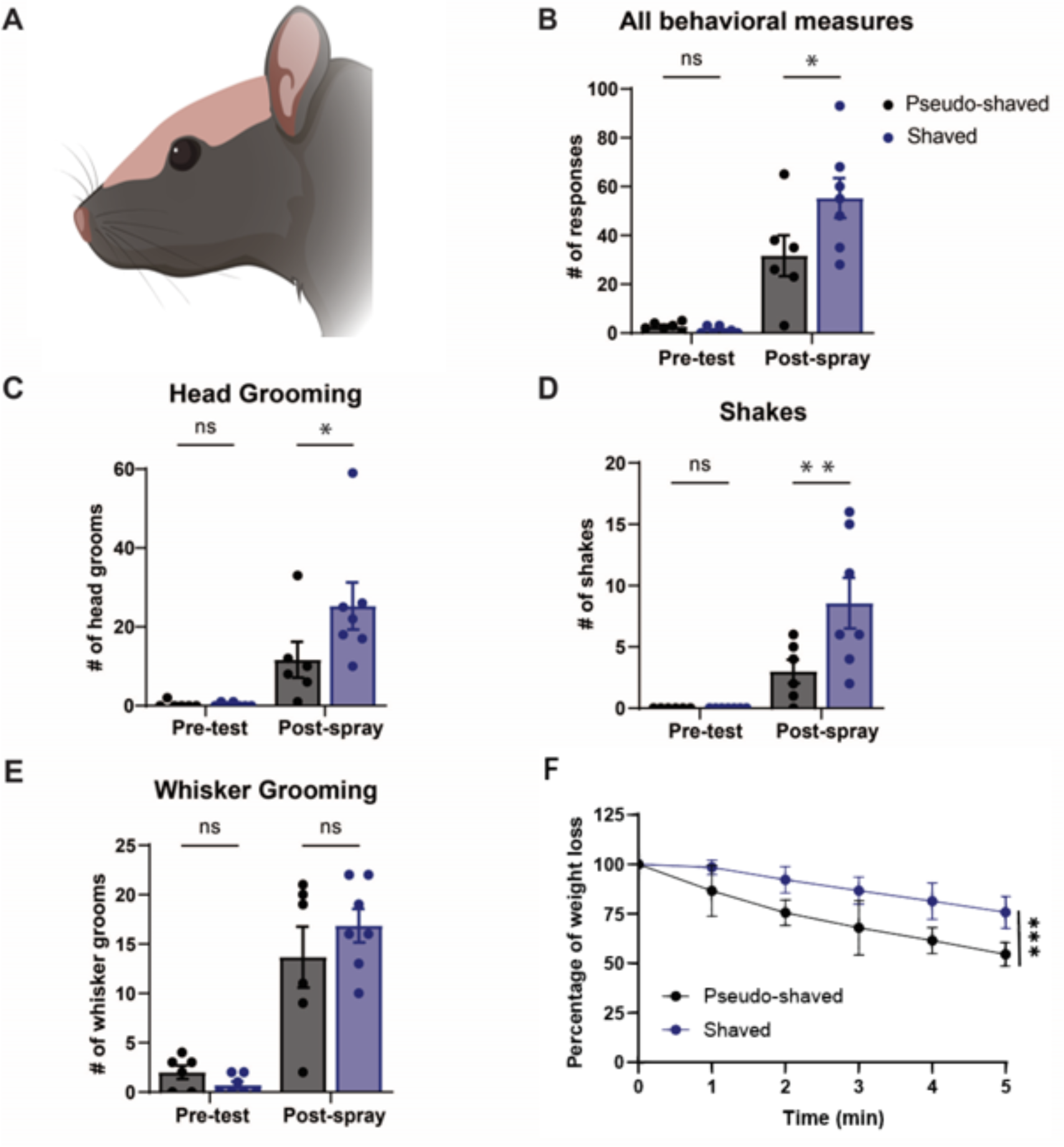
Head fur shaving enhances behavioral responses but slows drying after head spraying. **(A)** Illustration of a rat after dorsal head fur shaving. **(B)** Head-shaved rats perform more drying behaviors than pseudo-shaved rats after the head spray (*p= 0.0192). **(C)** Head-shaved rats head-groom more after the head-spray than pseudo-shaved rats (*p=0.0412). **(D)** Head-shaved rats engage in more shakes after the head spray than pseudo-shaved rats (**p= 0.0072). **(E)** Pseudo-shaved and head-shaved rats display no difference in whisker grooming post-head spray (p=0.3708). **(F)** Percentage of weight loss in control (n = 5) and shaved (n = 7) animals for 5 minutes. The total amount of water sprayed on was the weight difference before and after spraying the animal, the weight difference to the weight gain after spraying was measured every minute and plotted in % of the total weight gain. There is a significant difference in weight loss between rats that are shaved or not (***p=0.0005). Unshaved: black, n = 6, Shaved: blue, n = 7. Error bars indicate ± SEM. Data underlying this figure can be found at https://figshare.com/s/109204cbfff25915ce87

### Fur dries faster than shaved skin by exposing water on distal fur tips to air flow

We were surprised that head shaved animals dried slower than pseudo-shaved animals and wondered if we could study this phenomenon *in vitro*, i.e., through the analysis of fur patches. To validate our model, we applied thermal imaging and found that awake wet rats exhibited a similar temperature distribution as excised fur patches (Supplementary Figure S2). Specifically, fur apexes were 2-3°C colder than the surrounding fur both *in vivo* and in excised fur. Encouraged by this similarity we performed thermal imaging of shaved and non-shaved rat back fur (Figure 6A). In all five experiments, dry shaved and non-shaved rat back fur had a similar temperature to the environment. Wet shaved and non-shaved rat fur had a lower but similar temperature when airflow was blocked. However, when exposed to the ambient air, fur was colder than dry fur and wet shaved skin, and this effect was significant across experiments. We suggest this low temperature reflects the highly effective evaporative cooling of fur. When the fur/skin patch was flipped, we observed the opposite temperature distribution: wet skin was colder on the inside than wet fur. This observation suggests that wet fur poorly conducts heat. Furthermore, when we increased air flow by a small ventilator, the shaved skin temperature stayed the same on average, but fur patches warmed by on average 0.5°C. Given the presumed increase in evaporative cooling with more airflow, we take this as an indication that fur is very effectively warmed by air. Next, we checked evaporation from shaved and non-shaved rat back fur (Figure 6B). We found minimal evaporation from fur when air flow was blocked, but when we applied air flow, we observed significantly more evaporation from fur compared to shaved skin. To gain insight into the distribution of water in fur, we applied a black aqueous fluid to white rat back fur and observed a highly uneven fluid distribution, with the applied dye being primarily confined to the distal fur tips (Figure 6C). In a similar experiment, we visualized the distribution of X-ray dense KJ-solution in fur using microCT volume images. Again, the volume rendered images showed that fluid distribution is highly uneven, confined to distal fur tips and fluid filled apexes appearing to float above the skin (Figure 6D, left). After drying we found that KJ residues were preferentially distributed to distal hair tips (Figure 6D, right). We also observed that distal hair tips concentrate dye after drying; such effects were seen both for anillin blue/ orange G (Figure 6E) and for eosin (data not shown). To understand fluid-flow in wet fur, we rubbed water into rat back fur and then injected dye proximally into fur apexes (Figure 6F). Fluid is sucked rapidly into the apexes. Without exception dye spread distally very fast. There was very little dye movement proximally (in the direction of the skin) and dye movement occurred preferentially along hair bundles (Figure 6G). In conclusion, we propose that fur evaporates water more effectively than skin by drawing water into distal hair tips and exposing it to air flow. This process is how we suggest wet fur acts as a heat pump, extracting the heat required for drying by cooling the ambient air (Figure 6H).

**Figure 6.**
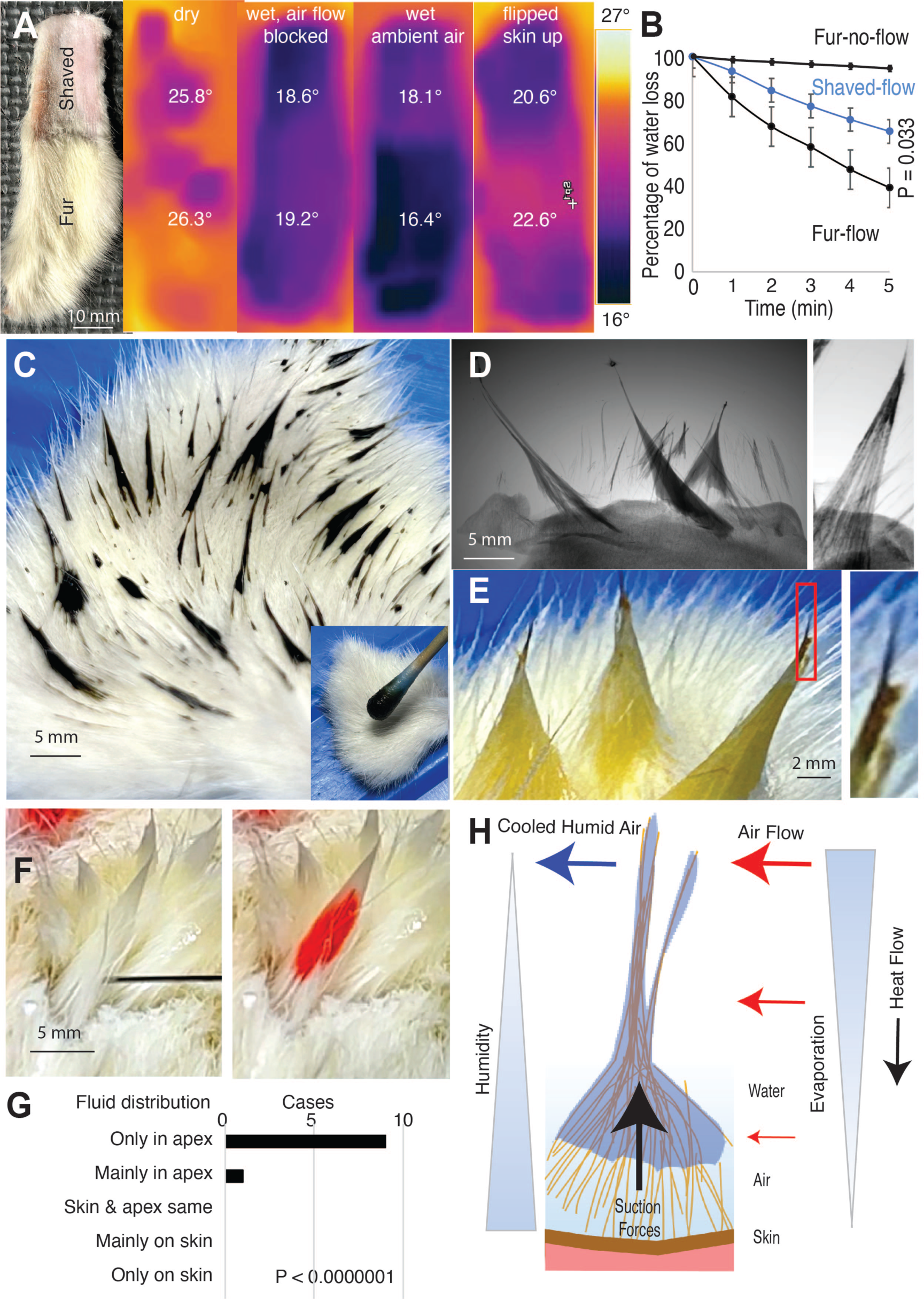
Fur dries better than skin by exposing water to air flow and hence wet fur might extract the heat required for drying by cooling ambient air. **(A)** Left, a partially shaved back skin patch was analyzed in thermal imaging. Fur and shaved skin have a similar temperature, when dry (left thermal image) and have similar lower temperature when wet and air flow is blocked by a petri dish (2^nd^ from left thermal image). The fur patch is colder than the shaved skin, however, when air flow is possible (3^rd^ from left thermal image: n = 5; fur avg -5.5° shaved avg -4.0° Celsius colder than ambient; ***p = 0.0002). Even though the fur is very cold, the inner skin below stays warmer than the inner skin below the shaved skin patch (4th from left thermal image, taken directly after flipping the fur patch: n = 5; fur skin avg -2.5° shaved skin avg -3.4° Celsius colder than ambient; *p = 0.03). **(B)** Percentage of weight loss in shaved skin (n = 5) and fur (n = 5) patches for 5 minutes. Air flow was either blocked (by closing the weighing chamber) or fur/ skin patches were exposed to air flow from a small desktop ventilator. The total amount of water sprayed on was the weight difference before and after spraying the patch, the weight difference to the weight gain after spraying was measured every minute and plotted in % of the total weight gain. There is a significant difference in weight loss between shaved skin and fur (*p= 0.033). **(C)** We rubbed (inset lower right) a black aqueous fluid (concentrated anillin blue/ orange G) into white fur to white rat back skin. Unexpectedly, the dye distribution is very uneven and confined to distal fur tips. **(D)** Left, in a similar experiment we visualized aqueous fluid distribution by applying X-ray dense 1% KJ-solution to fur and taking microCT volume images. We then thresholded images such that only fluid and skin were visible. The volume rendered images show fluid distribution is very uneven, confined to distal fur tips and fluid filled apexes float above the skin. Right, after drying the fur patch we find that KJ residues are preferentially distributed to distal hair tips. **(E)** Left, distal evaporation from fur tips was also suggested when we applied aqueous fluid (diluted anillin blue/ orange G) to white fur apexes and dried the fur for 30 minutes. After initial application of diluted anillin blue/ orange G fur apexes were entirely orange (not shown), but after drying the fur tips turned black (the color of concentrated anillin blue/ orange G). Right, black fur tip at higher magnification. **(F)** To understand the fluid flow in fur apexes we injected fur apexes proximally close to the skin (left image, the black injection needle) with red dye (right image; eosin red). Fur apexes were generated prior to the experiment by rubbing water in rat back fur. **(G)** Invariably injected fluid was pulled up in the apex and it reached only in one case the skin at all; note that all experiments were done with apex tips up (i.e., against the pull direction of gravity). We tested the significance of the effect with a binomial test against the null-hypothesis of equal probability outcomes. **(H)** The wet-fur-heat-pump-hypothesis. Based on a synopsis of our findings we suggest that wet fur dries very effectively by exposing water on distal fur tips to air flow. This is the way fur might act as a heat pump that extracts the heat required for evaporation from the ambient air rather than from the animal. Data underlying this figure can be found at https://figshare.com/s/c908dfebae47ae37b56d

## Discussion

### Summary

We found that spraying rats with water induces a host of changes to rat fur, including the formation of fur apexes, as well as alterations in behavior. Apex formation and behavioral wetness responses depend on the wakefulness of the animals and promote drying. Drying rates and behaviors evoked by water spraying differ across body parts. The facial skin is involved in water sensing and driving wetness responses. Shaving experiments point to a decisive correlation between fur and the drying process. Fur evaporates water more effectively than shaved skin. Suction forces move water into the distal hair tips and expose it to airflow. This mechanism appears to extract the heat required for drying by cooling the ambient air.

### Rats drying requires activity

A crucial insight from our study is that the drying process requires the animal to be active. Specifically, we found that awake water sprayed animals dry about four times faster than anesthetized water sprayed animals. This difference is quite remarkable, given that it is the same body that is wet in awake and anesthetized animals. We suggest that rats, and presumably other mammals, developed robust drying mechanisms to not suffer from hypothermia. We also noted that in terms of surface-to-volume considerations, integument wetness induced hypothermia is a much larger challenge for small homeothermic animals like rats than it is for humans [7].

### The wetness response

Water spraying leads to a host of changes in the sprayed rats, namely (i) behavioral changes, (ii) fur changes and (iii) other changes that were not measured in this study. The behavioral response component is very complex. The most common behavior evoked by spraying was whisker grooming, but we also observe increased grooming of body and head. Additionally, the animals increase scratching behaviors and shaking. Notably, the effect of shaking on water removal has been characterized and it is a very effective water removal strategy in soaking wet animals [3]. All of the behaviors (whisker grooming, head grooming, body grooming, scratching shaking) fall into the category of cleaning behaviors. However, in this study we sought to assess their role in water removal. We will discuss fur changes and other response components not measured here in detail below.

We propose that these behaviors and fur changes should be understood as part of a coherent behavioral program referred to as a wetness response, rather than as single behavioral elements. The body parts differ in their ability to evoke the wetness response, but the evoked behaviors are very similar; specifically, both head and back water spraying induce mainly whisker grooming. The impression of a fixed action pattern program is enhanced by the similarity of behavioral and fur changes seen in awake animals.

### Wetness sensing, the role of facial skin, is there a human wetness response?

Wetness sensing has been studied in both animals and humans [8]. The evidence suggests combination of mechanical and thermal inputs leads to wetness perception [5], with colder stimuli being perceived as more wet [9]. In line with such thermal effects, it has been suggested that Aδ afferent fibers play a role in human wetness perception [10]. A more general model of human wetness perception emphasizes the central integration of information from mechanical and thermal afferents [9].

Several lines of evidence suggest that the rat’s wetness response is powerfully activated from the rat’s dorsal facial skin: (i) We find that head spraying evokes strong wetness responses; (ii) Whisker trimming does not affect such responses evoked by head water spraying. (iii) Blocking the dorsal facial skin diminishes wetness responses. (iv) Exposing the dorsal facial skin to wetness by shaving potentiates wetness responses. Previous studies suggest that there is no special wetness sensitivity of the human face [11].

It is clear, however, that the human facial skin also mediates unique and very powerful responses to water. Such responses to water immersion of the human face were discovered in the context of research on diving and have been referred to as the human ‘diving response’ [12,13]. Specifically, immersion of the human face in water evokes powerful cardiovascular responses such as bradycardia and peripheral vasoconstriction [14,15] and such responses are not seen for body immersions excluding the face [16]. Most interestingly, many if not all aspects of the so-called diving response are blocked by thermal adaptation of facial cold receptors [17]. The strong cardiovascular responses to face wetting in humans suggest that it would be worthwhile to assess the cardiovascular effects that occur simultaneously with the rat wetness response. It is also obvious that many aspects of the so-called human diving response (dependence on facial thermal inputs, peripheral vasoconstriction) might also function as a defensive thermal reaction. Given that getting wet is much more common than going for a dive, we wonder if parts of the human diving response may have evolved an energy-preserving wetness response.

### A decisive role of fur in drying

Our data indicate a crucial role of fur in rat drying. Specifically, we find that head shaved rats dry poorly, even though they perform intense grooming and shaking behaviors. Our *in vitro* data show that fur dries more effectively than shaved skin, and that this effect is strongly air flow dependent. We reason that the very slow drying of anesthetized animals can be attributed to their lack of movement and the resulting absence of airflow. Fur changes, supported by active mechanisms such as piloerection [18], appear to aid in apex formation and might thereby accelerate drying. Similarly, shivering may strongly increase air flow around the fur. Several lines of evidence suggest that evaporation predominantly occurs from distal fur tips. First, apexes and their tips are colder than the surrounding fur, both *in vivo* and *in vitro*. Second, apexes and their tips concentrate dyes during drying *in vitro.* Third, apexes and their tips concentrate KJ-salt during drying *in vitro*.

### The wet fur heat pump hypothesis

Our *in vitro* data provide a good intuition for how fur promotes drying. Wet fur exposed to air flow gets very cold, likely due to evaporative cooling. However, the inner skin side of wet fur stays warmer than the inner skin side of wet shaved skin. This occurs because evaporation appears to be happening preferentially on the very distal hair, and suction forces push the water distally. These suction forces appear to act along hair bundles, and we wonder if they arise from capillary forces associated with hair properties. The tepee-like architecture of fur apexes, with higher hair density distally than proximally, might also contribute to these forces. Evaporation increases with air flow and decreases with humidity [19]. Accordingly, we suggest that the strong distal airflow and the distally lower humidity explain the preferential evaporation from the very distal hair (Figure 6H). Furthermore, we suggest that, as a whole, the exposure of water to air flow on distal hair tips might act as a heat pump that extracts the heat required to evaporate the water by cooling the ambient air. Several factors might contribute to the effectiveness of this heat pump mechanism. First, because evaporation occurs distal from skin, i.e., the distances for conducting the low temperature from evaporative cooling to the skin are long. Second, water does not fully saturate the fur and therefore skin is isolated from water-containing apexes by air bags. Third, distal hair tips are long and thin, resulting in poor vertical conduction of low temperature to the animal but efficient heat exchange with the surrounding air. Fourth, distal airflow is strong (both as a result of ambient air flow and because of the locomotion of the animal), but proximal airflow is weak (because of the air trapping properties of fur). With such a heat pump mechanism, the heat required for drying comes largely from the ambient air, i.e., for free.

### Hairless mammals: did fur evolve for drying?

Hair is a defining characteristic (the common synapomorphy) of mammals. The lack of fur in poikilotherm animals has long suggested a relation of fur and thermoregulation. To better understand the relation between fur and thermoregulation, it might be worthwhile to consider fur loss in mammals. Fur loss is very rare in wild mammals, even though there are numerous viable hairless mutants in dogs, cats, rats, guinea pigs etc. This observation suggests that hairless mutants are constantly eliminated by selection. The only larger group of mammals that has fully lost fur is the cetaceans. These mammals are fully aquatic and hence they never dry. Some semi-aquatic mammals, such as the hippopotamus, have lost fur, but most semi-aquatic mammals (otters, seals) retained their fur and they regularly dry. Very few land mammals lost fur. These include the northern babirusa, a large pig species, which lives in a warm constant temperature environment. Another furless species is the naked mole rat, which lives a subterranean lifestyle and presumably is only rarely or never exposed to rain and drying. Extant elephant species lost hair and at least African elephants make big migratory efforts to avoid rain (and drying?). Humans have also lost fur, and there is no other species that goes to same length to avoid rain. Human efforts to avoid rain include caves, coats, huts, houses and holidays in the sun. We conclude that many furless mammals dry either never or only rarely (cetaceans, the naked mole rats) or undertake major efforts to avoid rain (elephants, humans).

### Conclusion

In response to water spraying rats actively promote drying by fur changes and a complex program of putative water removal behaviors. Fur shaving decisively delays drying and fur promotes evaporation by sucking water into distal hair apexes and exposing it to air flow. We hypothesize that this is the way fur acts as a heat pump and extracts the heat required for drying by cooling the ambient air.

## Materials and methods

### Animals

Long–Evans rats (P21–P32, n = 35) were used in these experiments. All experiments complied with regulations on animal welfare and were approved according to international law for animal welfare and approved by Woods Hole, USA (22-09E and 23-09C), and the Institutional Animal Care and Use Committee (IACUC).

### Water spraying response behavior

Long–Evans rats (P21–P32) were separated from littermates before behavioral testing. Behavioral videos were recorded with a Logitech camera (C920x HD Pro, 1080p/30fps) while rats were placed in a glass fish tank or plastic beaker placed upon a scale (Adventurer Analytical, Ohaus).

Rain stimuli consisted of spraying rats 5 times focally on the specified body parts after a baseline period. Experimenters then scored behavioral responses and confirmed post-hoc with video analysis (Behavioral Observation Research Interactive Software). In weight loss experiments, rats were placed inside the plastic beaker that was positioned on the center of a scale. The weight of the beaker was accounted for in the baseline period, and the rat’s weight was assessed before and after spraying and every minute thereafter.

### Whisker trimming or lidocaine/Ringer injections

Whisker trimming or lidocaine/ringer injections were performed in gently restrained animals under stereoscopic magnification and illumination. Injections were performed subcutaneously and directed to two spots on the head. Head fur was trimmed with an electric razor. Sham trimming/injection procedures were implemented to control for the stress of shaving. In such sham procedures, animals were gently restrained, positioned under the microscope, and scissors and razors were brought close to the animal’s face.

### In vitro analysis of fur and skin patches

To obtained rat fur we skinned dead rats (n =5). Large fur patches were easily obtained from back or belly skin and all the *in vitro* analysis focused on back skin. Patches were shaved and sprayed in the same way as described for awake rats. Fur apexes were generated by rubbing wet fur with a wet Q-tip against the laying direction of the fur.

We rubbed either KJ-solution, anillin blue/ orange G dyes into fur. Fur apexes were injected with eosin solution proximal to the skin under visual control under a magnifying glass. All injections were videotaped (in slomo modus) and photographed before and after injection using an iphone 14.

### MicroCT Imaging

We used diffusible iodine-based contrast enhanced computed tomography (diceCT) to analyze and visualize the distribution of 1% KJ solution in fur. Computed tomography scans were obtained by YXLON FF20 CT system (YXLON International GmbH, Hamburg, Germany; RRID: SCR_020903). Scans were performed with an isotropic voxel size of 10.6 µm. Images were visualized and volume rendered version of the Amira (Amira ZIB Edition 2022, Zuse Institute Berlin, Germany) and dragonfly software.

### Thermal Imaging

Thermal imaging was done with a Flir One Gen C camera (Flir systems). Digital and thermal images were processed with the Flir Tools software (Flir Systems). Thermal imaging was performed on dry rats, water sprayed rats and excised fur and skin patches. Air flow was blocked by putting petri dishes over fur specimen.

### Statistics

Most of our dataset did not satisfy normality criteria, so we applied nonparametric statistics. We analyzed data from binomial distributions with χ^2^ and Fisher’s exact test. Mann–Whitney, Wilcoxon, or Kruskal–Wallis tests were employed to analyze 2 unpaired groups, 2 paired groups, or more than 2 unpaired groups, respectively. One-way Anova and post hoc Tukey analysis were used for Figure 1F-I. Additionally, Two-way repeated measures ANOVA and post hoc Šidák analysis were used for Figure 4B, D and Figure 5B-E. We used a linear regression analysis for Figure 5F. Data was expressed as the mean ± the standard error of the mean (SEM). We only report differences which were significant or relevant to the experiment. In all cases, *p* < 0.05 was the statistical threshold. The analyses were done using Prism or MATLAB (MathWorks, Natick, Massachusetts, USA).

## Acknowledgements

This work was supported by the Marine Biological Laboratory, a training grant from the NIMH (R25MH059472), Humboldt Universität zu Berlin, the Bernstein Center for Computational Neuroscience Berlin, the German federal ministry of education and research. Ann Clemens is supported by the Simons Initiative for the Developing Brain, the University of Edinburgh and a Simons Edinburgh Scientific Academic Track (Simons-ESAT) fellowship. LZ was supported by Patricia A. Case Endowed Scholarship; ATA and PNM were supported by a Marine Biology Laboratory Award; MAD was supported by The Grass Foundation to attend the Neural Systems & Behavior Course (NS&B). We thank Alberto Pereda, Stephanie White, Rosalie Maltby, Rose Holzhauer, and the Neural Systems & Behavior folks; this is the way.

## Supplementary Material

### Supplementary Figure 1

**Fig. S1.**
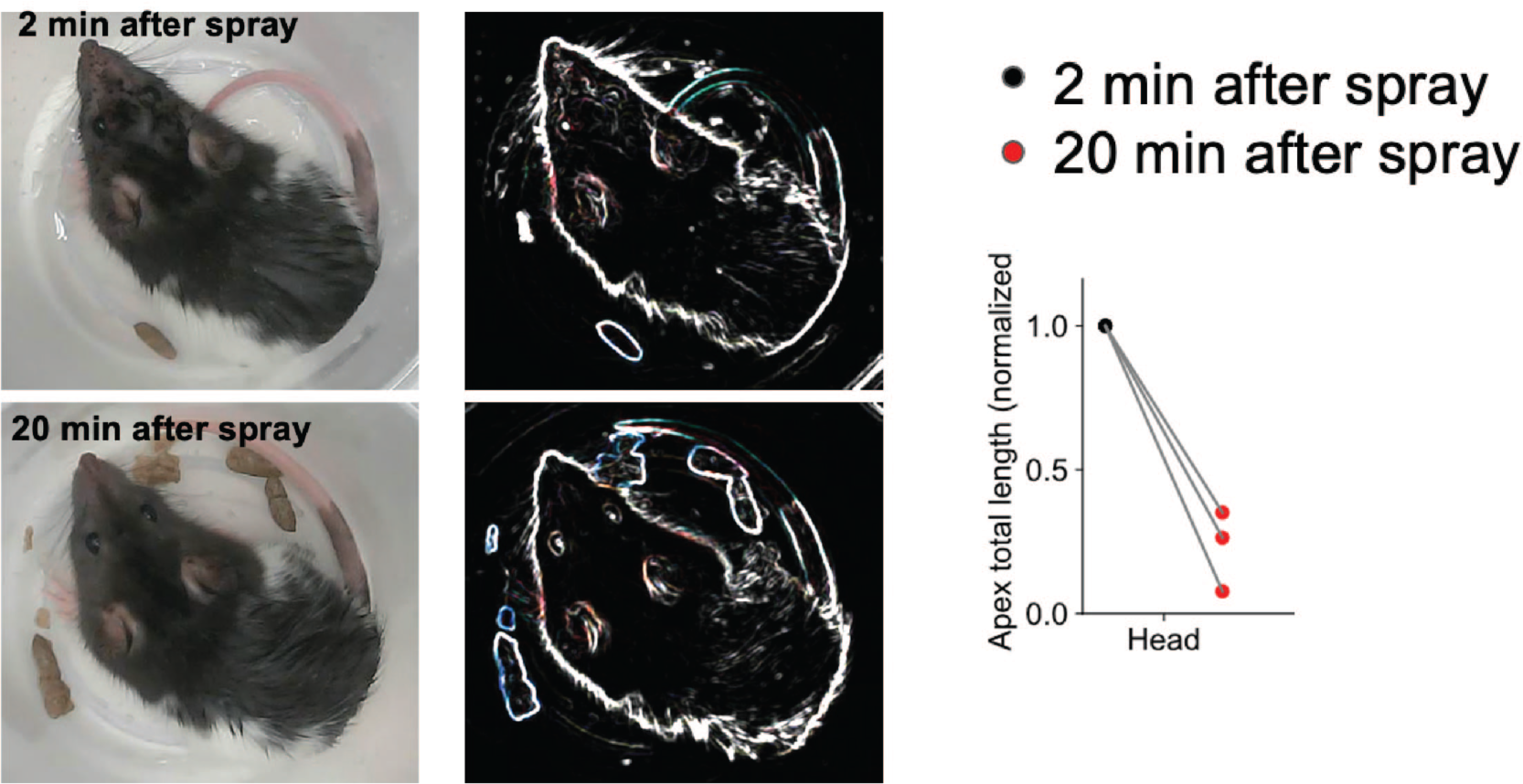
Quantification of the appearance and disappearance of fur apexes in the drying process. **(A – B)** Photographs of a rat showing 2 min **(A)** and 20 min **(B)** after water spraying on the head and body. **(A’ – B’)** Variance filtered **(A)** and **(B)** exemplifying the results of edge detection and apex locations on the head (yellow arrow). **(C)** We quantified apexes (elongated tapering hair clumps) as the normalized total length of edges (visible as white lines after edge detection) at the 2 min and 20 min after spray (n = 3 animals); the disappearance of apexes as a result of drying is well visible. Data underlying this figure can be found at https://figshare.com/s/9985a434fd669361cabe

### Supplementary Figure 2

**Fig. S2.**
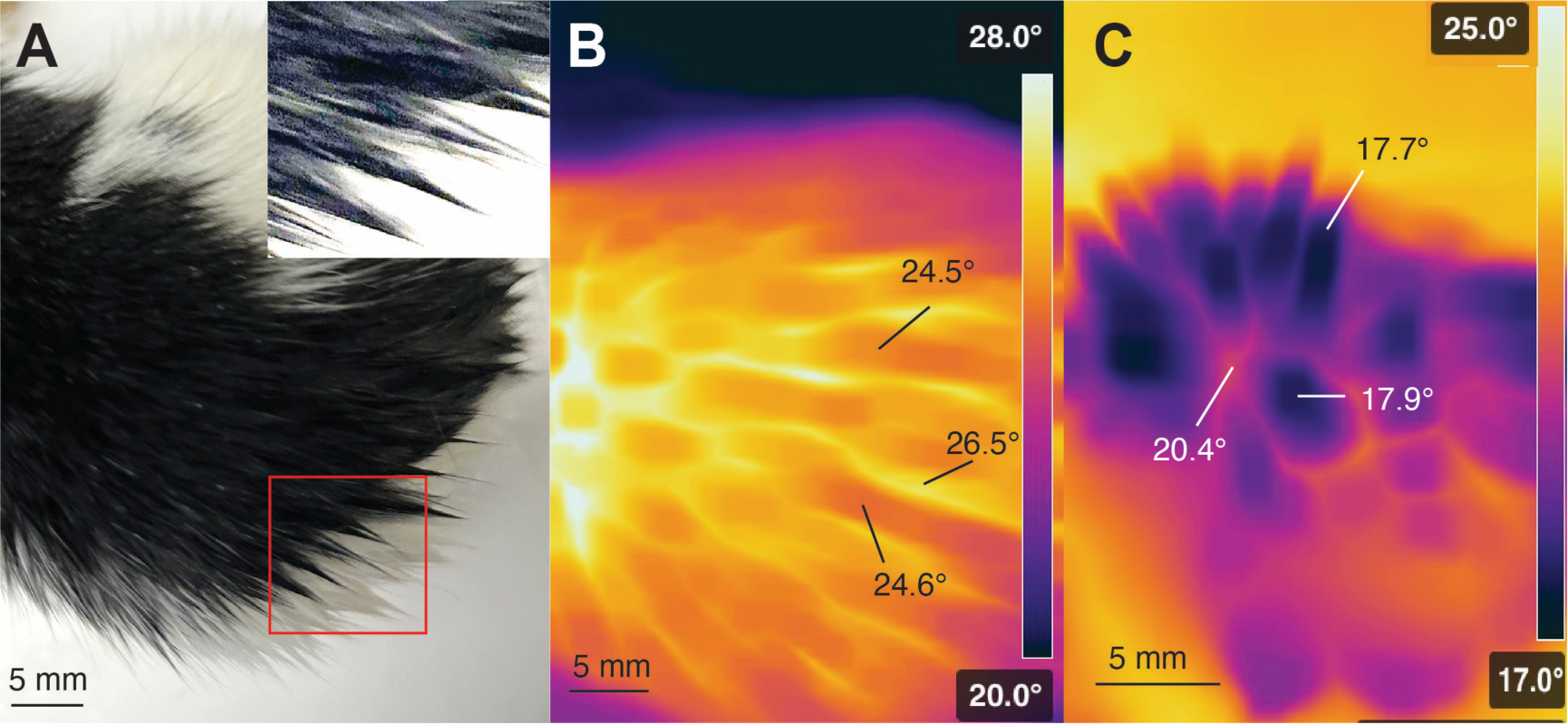
Photography and thermal imaging of wet fur apexes *in vivo* and *in vitro*. **(A)** Photograph of the back fur of a wet rat after water spraying as described before. Upper right inset shows fur apexes. **(B)** Thermal imaging of the wet back of the same animal. A regular pattern of colder fur apexes is seen; apexes are 2-3° Celsius colder than in between fur areas. **(C)** Thermal imaging of excised back fur. Apexes were generated by rubbing water into back fur against the laying direction of the fur. As *in vivo* (B) a pattern of colder fur apexes is seen; apexes are 2-3° Celsius colder than in between apex fur areas.

